# In vivo and in vitro human gene essentiality estimations capture contrasting functional constraints

**DOI:** 10.1101/814855

**Authors:** JL Caldu-Primo, JA Verduzco-Martínez, ER Alvarez-Buylla, J Davila-Velderrain

## Abstract

Gene essentiality estimation is a popular empirical approach to link genotypes to phenotypes. In humans, essentiality is estimated based on loss-of-function (LoF) mutation intolerance, either from population exome sequencing (in vivo) data or CRISPR-based in vitro perturbation experiments. Both approaches identify genes presumed to have strong detrimental consequences on the organism upon mutation. Are these genes functionally distinct and constrained by having key roles? Do in vivo and in vitro estimations equally recover these constraints? To address these questions, here we integrated disparate genome-scale datasets and compared structural, functional, and evolutionary features of essential genes versus genes with extremely high mutational tolerance and proteome expectation. We found that essentiality estimates do recover functional constraints. However, the organismal or cellular context of estimation leads to functionally contrasting properties underlying the constraint. Our results suggest that depletion of LoF mutations in human populations effectively captures developmental, organismal-level functional constraints not experimentally accessible through CRISPR-based screens. Finally, we identify a new set of genes (*OrgEssential*), which are intolerant of LoF mutation in vivo but highly tolerant in vitro. These genes drive observed functional constraint differences and have an unexpected preference for nervous system expression.

## Introduction

Understanding the patterns and phenotypic consequences of genotypic alterations is a fundamental problem in evolution and development ^1–4^. A popular empirical approach to link genotypes to phenotypes is by estimating the degree of *essentiality* of a gene. A gene is considered “essential” if it is required to sustain life in cells or whole organisms, and this requirement is often estimated by experimental perturbations ^5,6^. Thus, the study of essential genes was originally conducted on prokaryotes, due to their accessibility to genetic manipulation. More recently, however, gene essentiality has been estimated in multicellular eukaryotes, including mammals ^7^. Despite the absolute character of the “essential” gene denomination, data from multiple studies in model organisms have shown strong context dependency: genes are required for survival or not depending on environmental conditions and developmental stages ^5,8,9^.

Sequencing technologies and gene editing techniques enabled the estimation of gene essentiality in humans ^6^. The problem has been addressed following two approaches. On one hand, systematic testing of gene silencing effects on human cell cultures identifies genes that affect cell viability or optimal fitness upon perturbation ^10–13^. On the other hand, population-level statistical estimates of unexpected mutational depletion identifies genes presumed to be subjected to functional constraints^14^. Both approaches aim at ranking genes according to their effect on the organism (or cell) upon loss-of-function mutations. However, given the context dependency of gene essentiality, and the differences in the organizational level at which the effects of genotypic changes are assessed, the parallels of the two types of essentiality approximations are unclear.

*In vitro* screens of mutation tolerance identify genes with an immediate effect on cell proliferation and viability; consequently, the corresponding essentiality estimates depend on the specific cell line and culture conditions being tested. In addition, cell culture experiments do not capture developmental and functional constraints intrinsic to the organism. *In vitro* estimation of gene essentiality is thus inevitably tailored to cell viability and/or fitness. On the other hand, “*in vivo*” measures of mutational tolerance estimated from population-level genetic variation score genes according to the prevalence/depletion of loss-of-function (LoF) mutations. Genes showing mutational constraint are assumed to be consistent with a scenario where purifying selection filters out protein altering mutations with detrimental effects, thus eluding fixation within the population. In this sense, *in vivo* estimates of mutational tolerance are considered a proxy for the effect of mutations on organismal fitness. Such effect, in turn, mirrors to some extent the notion of essentiality in the context of population dynamics ^5^. Both types of estimations (*in vitro* and *in vivo*) have been discussed within the context of human gene essentially, nonetheless ^6^. Hereafter we use the terms cellular viability (CV) and organismal fitness (OF) to refer to the context at which human gene essentiality is estimated by means of either *in vitro* perturbation-based or *in vivo* population-based mutation tolerance estimation, respectively.

Notably, in both the CV and the OF context, a subset of mutational intolerant genes has been observed, leading to the idea of defining an “essential genome” containing genes that do not tolerate mutations, and a “dispensable genome” including mutation-tolerant genes ^6,14^. Intolerant genes (essential) are commonly of interest due to their potential detrimental effect on phenotype and disease association; however, highly tolerant genes (non-essential) might be relevant for evolvability, due to the plasticity they confer to the system at longer time-scales -- for example, as sources of cryptic genetic variation ^4^ or possible editable links that integrate subsystems ^15^. Hereafter we will use the terms *tolerant* and *intolerant* to refer to human essential or non-essential genes as estimated by the degree of tolerance to LoF mutations.

Despite the potential functional relevance of tolerant and intolerant genes, an understanding of the molecular determinants that discriminate the two groups has been only partially explored for humans ^6^. Moreover, an understanding of the dependency of molecular determinants of gene essentiality on the differences between the operational context of estimation (CV, *in-vitro* vs. OF, *in-vivo*) is lacking. To address these problems, here we systematically defined groups of human tolerant and intolerant genes and performed an integrative and comparative analysis of the structural, functional, and evolutionary features associated with gene essentiality. We analyzed the particularities and commonalities between genes that show extreme (in)tolerance to LoF mutation in a given context: CV, OF, or both (Figure 1).

**Figure 1.**
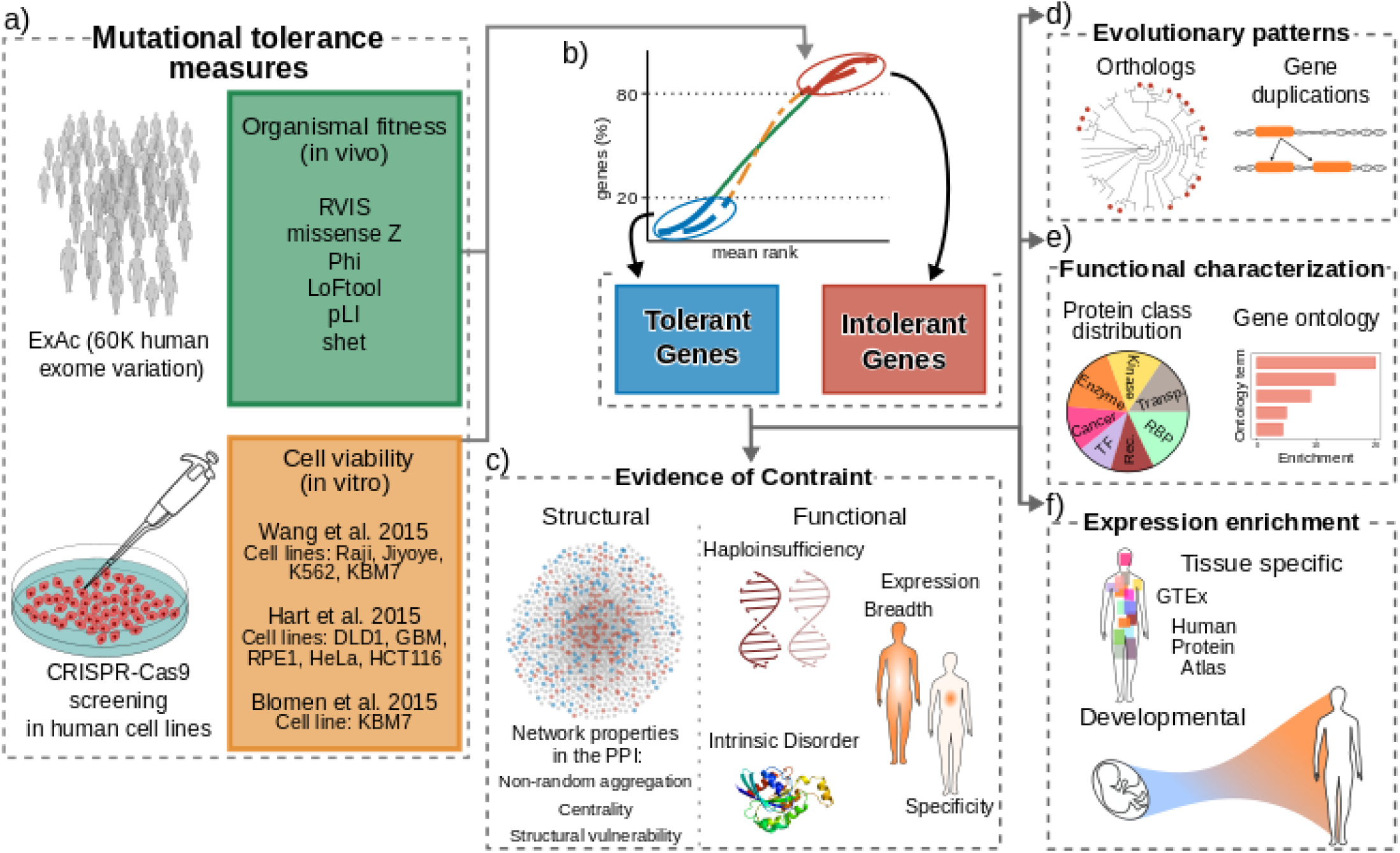
Overview. **a**, Mutational tolerance scores used to categorize human (in)tolerant genes. **b**, Consensus mutational tolerance score derivation (mean rank distribution) and corresponding (in)tolerant gene sets. **c-f**, Features considered as potential determinants of mutational constraint and gene essentiality, including structural and functional features **(c)**, evolutionary **(d)**, protein functional characterization **(e)**, and expression enrichment **(f)**.

## Results

### Context-dependent mutational tolerance categorises human genes

To define genes that show extreme (in)tolerance to detrimental mutation, as estimated from patterns on mutational depletion in exome sequencing data (OF) or CRISPR-based cell culture perturbation experiments (CV) data, we first calculated for each gene and context a consensus tolerance score measured by the average rank across a total of 6 and 10 previously proposed tolerance measures for OF or CV, respectively. A high correlation among pairs of individual measures within each context (OF or CV) justifies the definition of the proposed consensus score (Figure 2a). We then defined a set of mutation-intolerant (tolerant) genes based on the distribution of the computed scores, using the 80th percentile of the distribution as arbitrary cut-off value. This choice approximates the previously proposed cut-off of pLI>0.9 used in ^14^ when introducing the widely used LoF-intolerance metric, pLI. In addition, we defined a contrasting, similar-sized set of mutation-tolerant (intolerant) genes by selecting the 20% bottom ranked genes of the consensus score distribution (Figure 2b). The size of the resulting gene sets are 3,028 genes for both intolerant and tolerant OF groups and 3,139 genes for both intolerant and tolerant CV groups. Exploring the intersection between gene sets, we identified 714 tolerant and 771 intolerant genes with consistent tolerance behavior across contexts. In contrast, we identified 918 inconsistent genes, whose tolerance behavior depends on the context (OF/CV) (Figure 2c).

**Figure 2.**
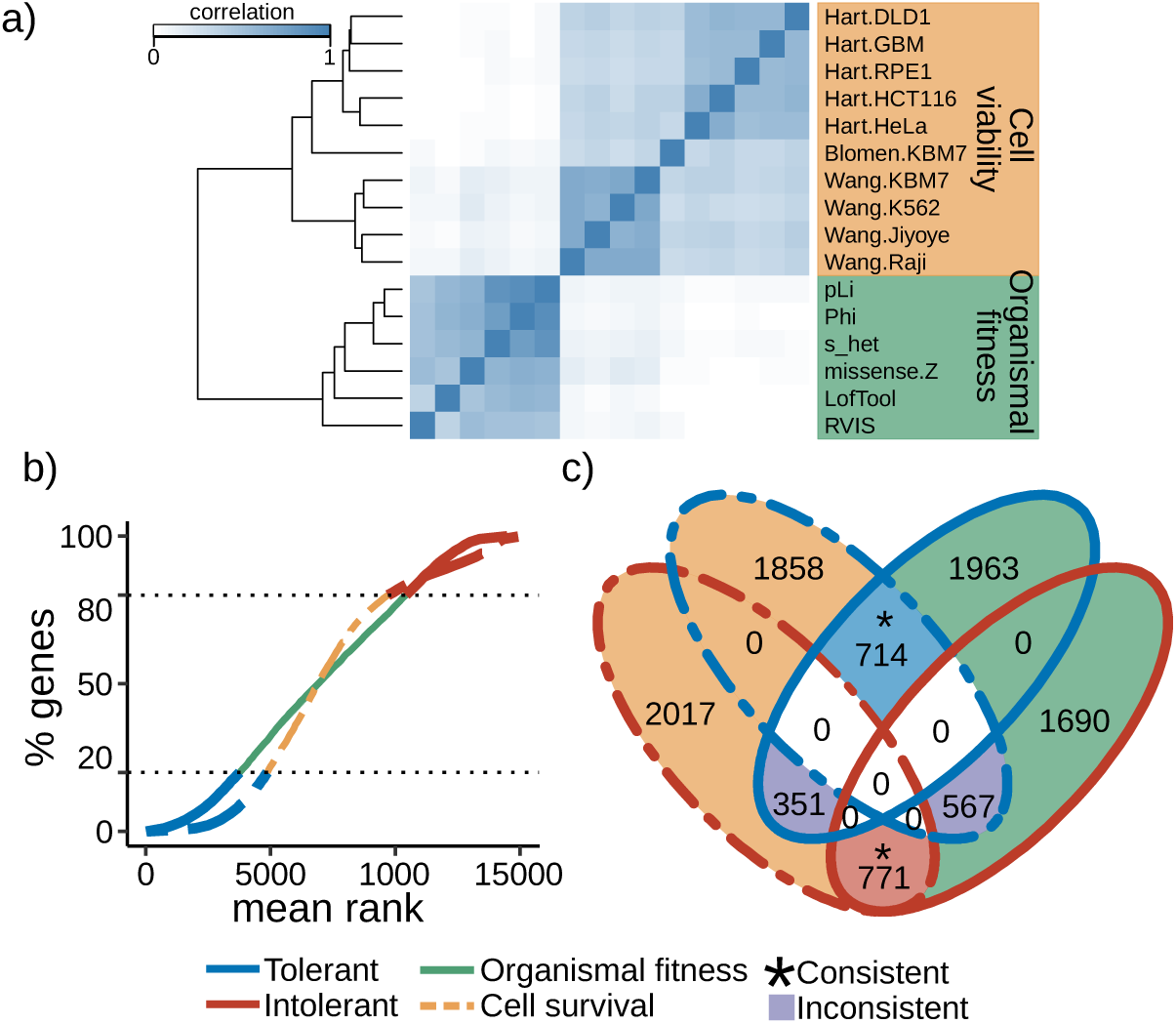
Mutational tolerant and intolerant groups definition. **a**, Correlation plot of mutational tolerance measures. **b)** Mutational tolerance measures mean rank distribution. Tolerant (blue) and intolerant (red) gene sets are defined as the bottom and top 20%, respectively. **c**, Venn diagram of the defined gene sets. Group intersections are highlighted to represent tolerant (blue) and intolerant (red) genes found consistently in both OF and CV groups, and inconsistent gene (violet) found in contradictory groups depending on the context.

### Structural and functional constraints predict mutational tolerance classes

We next tested whether the different gene groups are distinctively associated with structural and functional molecular network properties. Following previous studies that point to a central role of essential genes in the interactome ^16–19^, we first asked if (in)tolerant genes have contrasting positions in the interactome and whether such pattern is consistent in genes affecting both organismal and cellular fitness. From a network perspective, we hypothesized that (1) a core constrained neighborhood exists within the human interactome, which is separated from a more peripheral, scattered layer formed by mutationally tolerant genes; and (2) that, given the central role of the intolerant neighborhood, perturbations affecting the corresponding genes are more likely to confer structural fragility to the entire system.

By measuring network features associated with node centrality and aggregation (Figure 3a, see Methods), we confirmed that intolerant genes estimated by either effect on OF or CV, both have a significantly higher aggregation and centrality in the interactome, while tolerant genes consistently show the opposite behavior: loose aggregation and peripheral positioning (Figure 3c, Supplementary Figure 1). To test the vulnerability of the interactome to perturbations targeting tolerant or intolerant genes, we analyzed the network’s behavior as a function of the progressive removal of nodes in decreasing order of mutational intolerance score (Methods). This analysis further confirmed that there is a strong association between mutational patterns and the structural properties of the interactome, revealing that intolerant gene removal produces a higher structural damage than random node removal (Figure 3b).

**Figure 3.**
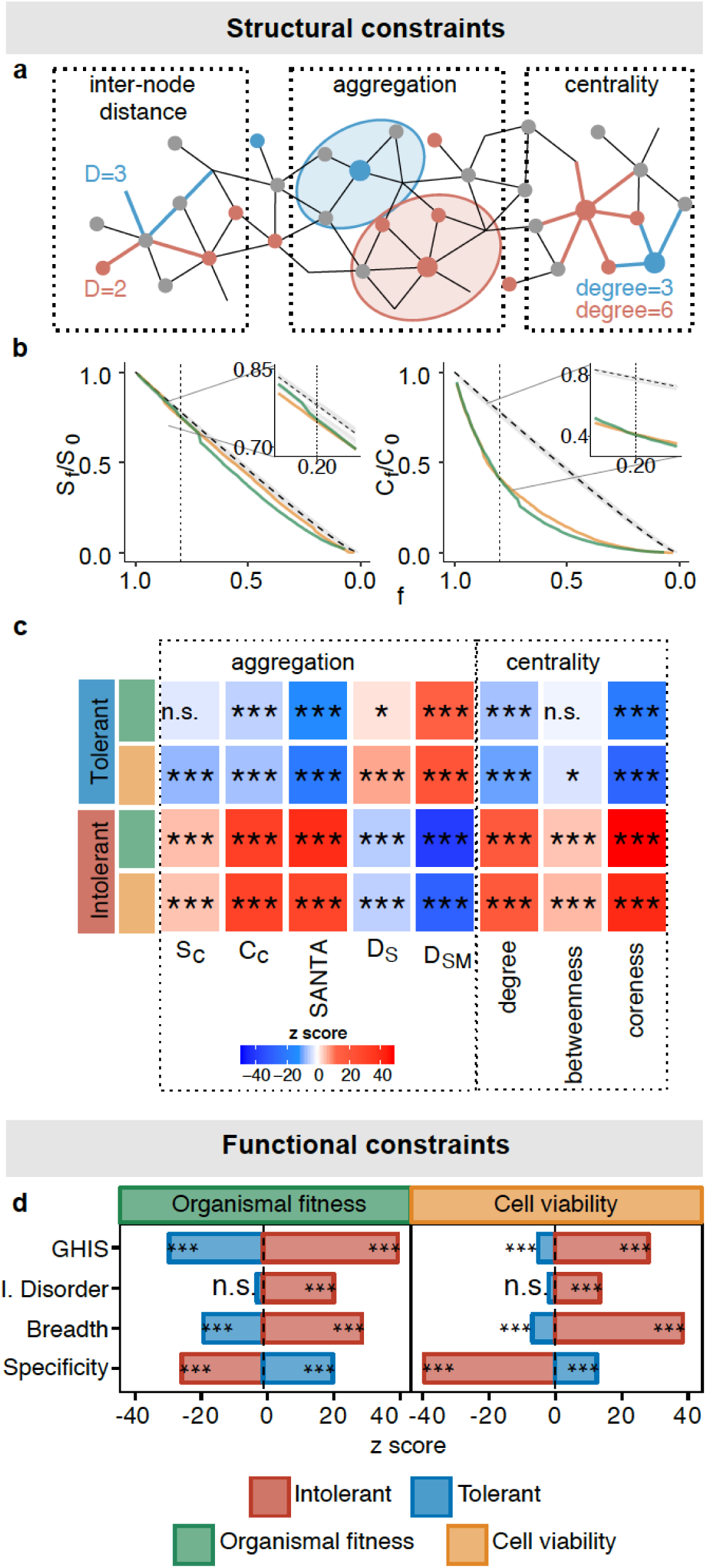
Structural and functional constraints. **a**, Network features measured in the PPI. **b**, Structural robustness of the PPI after random removal of nodes (gray lines) and directed removal of genes ranked by mutational intolerance score (inset shows the pattern around the 20% of nodes removal, corresponding to the intolerant gene sets). **c**, Deviation (z score) of every network feature. (significance: *p* < 0.05 = n.s., 0.001<*p*<0.05 = *, 0.0001<*p*<0.001 = **, *p*<0.0001 = ***).

The previous data suggests that global structural properties of the interactome are related to mutational tolerance classes. We reasoned that there might also be molecular properties suggestive of functional constraint that similarly discriminate tolerance groups. By analysing the degree of estimated haploinsufficiency (GHIS), protein intrinsic disorder (ID), expression breadth, and specificity; we similarly found contrasting patterns between tolerant and intolerant genes. Intolerant genes are more likely to be haploinsufficient, to have more intrinsic disorder in protein structure, and to be more broadly expressed across tissues; while tolerant genes show exact opposite behavior (Figure 3d, Supplementary Figure 1). Together, our results confirm that mutationally tolerant and intolerant genes can be consistently discriminated by features indicative of structural and functional constraints, irrespective of the context in which tolerance is estimated (OF or CV).

### Evolutionary history of tolerance gene classes

Gene essentiality and interactome centrality have been previously related to evolutionary conservation, with a tendency for topologically central and essential genes to be conserved (evolutionarily old) ^20,21^. We studied whether the tolerance gene groups identified here similarly have contrasting evolutionary conservation patterns and whether associations are consistent in genes affecting organismal or cellular fitness. We analyzed two features of evolutionary conservation: orthology and paralogy.

First, we evaluated the degree of gene conservation by calculating a gene conservation index (CI) that measures the proportion of species in which a human gene has a one-to-one ortholog (Figure 4a). We considered a total of 187 species from 7 taxonomic groups (Archaea, Bacteria, Protozoa, Fungi, Plants, Invertebrates, Vertebrates) (Methods). Intolerant genes are significantly more conserved than tolerant genes in both OF and CV contexts (Figure 4b, Supp. Fig. 2). Notably, however, the conservation of intolerant genes that affect cell viability is considerably higher than that of genes affecting organismal fitness. To further explore the difference in conservation, we calculated the proportion of genes with a one-to-one ortholog in each taxonomic group, for every tolerance group (Figure 4c). This analysis revealed a clear difference between CV and OF gene groups. In particular, CV intolerant genes are significantly more represented in every taxonomic group except for Vertebrates, while the behavior of OF intolerant genes does not deviate from random expectations, presenting only a marginal enrichment among Plants and Fungi and depletion in Bacteria (Figure 4c). This indicates a deep conservation of intolerant genes affecting cellular viability in humans and possibly reflecting the relevance of these genes in core cell-autonomous functions.

**Figure 4.**
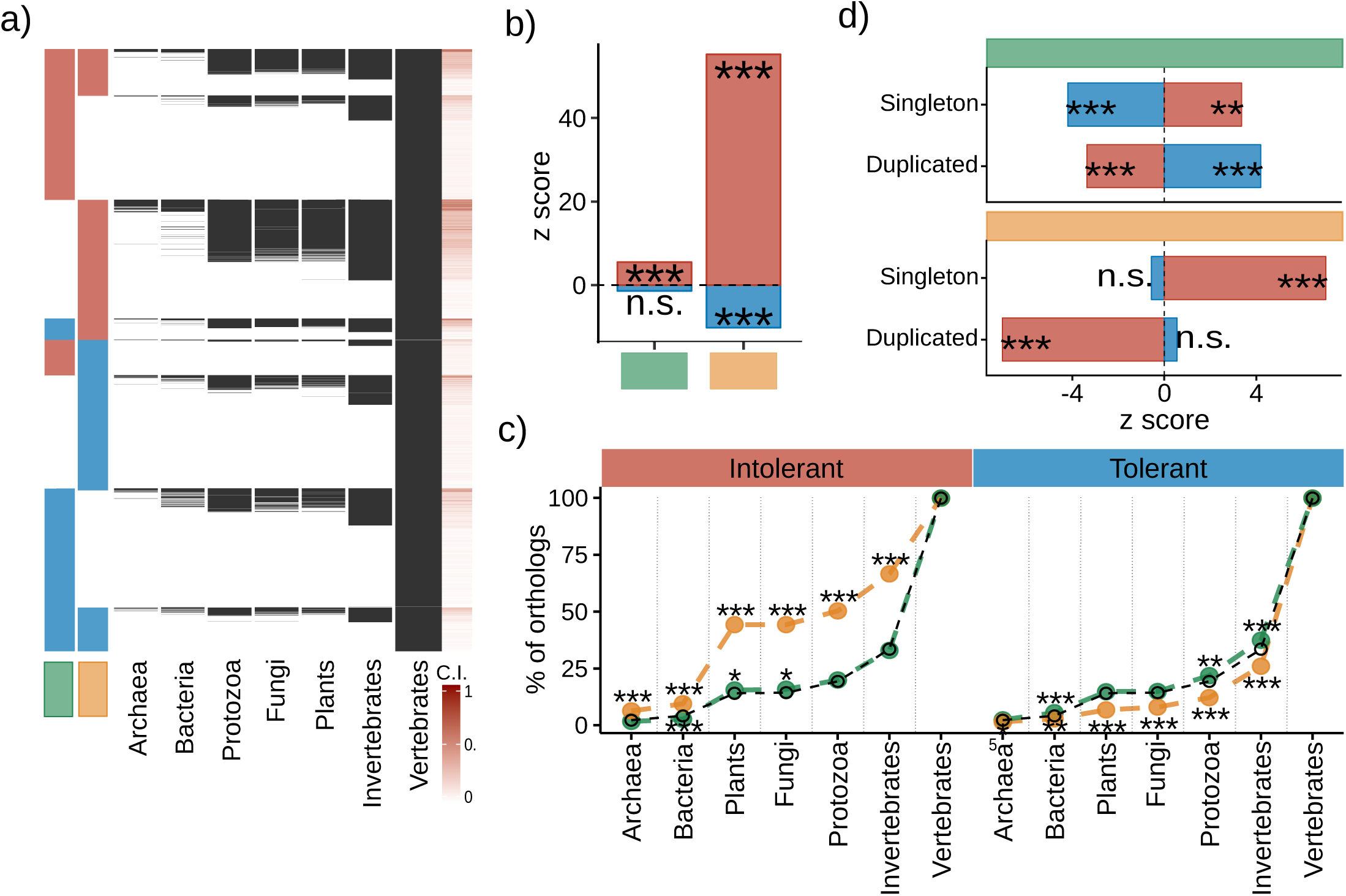
Genes evolutionary features. **a**, Presence of gene orthologs among taxa, each row shows the presence (black) of an ortholog in the given taxon. **b**, Deviation of mean gene C.I. **c**, Percent of genes with an ortholog in each taxon (random expectation is shown in black). d) Deviation in the number of singleton and duplicated genes per set. (significance: *p* < 0.05 = n.s., 0.001<*p*<0.05 = *, 0.0001<*p*<0.001 = **, *p*<0.0001 = ***).

We next analysed the association between tolerance groups and copy number variation, considering the number of gene duplication events represented in each gene group. The number of duplication events is consistently depleted among intolerant genes in both OF and CV, suggesting further evolutionary constraint in mutational events for genes that are also intolerant to deleterious point coding mutations. In contrast, tolerant genes show overrepresentation of duplication events, but only in the OF context, which is consistent with a scenario of relaxed selection pressure in these genes (Figure 5d). The evolutionary pattern of reduced duplication events in intolerant genes is also consistent with a reduction in gene family size distribution for intolerant relative to tolerant genes (Supp. Fig. 2). Together, these results confirm that there is a marked difference in the evolutionary history of tolerant vs intolerant genes. They also uncover clear differences in genes that affect cell variability or organismal fitness.

**Figure 5.**
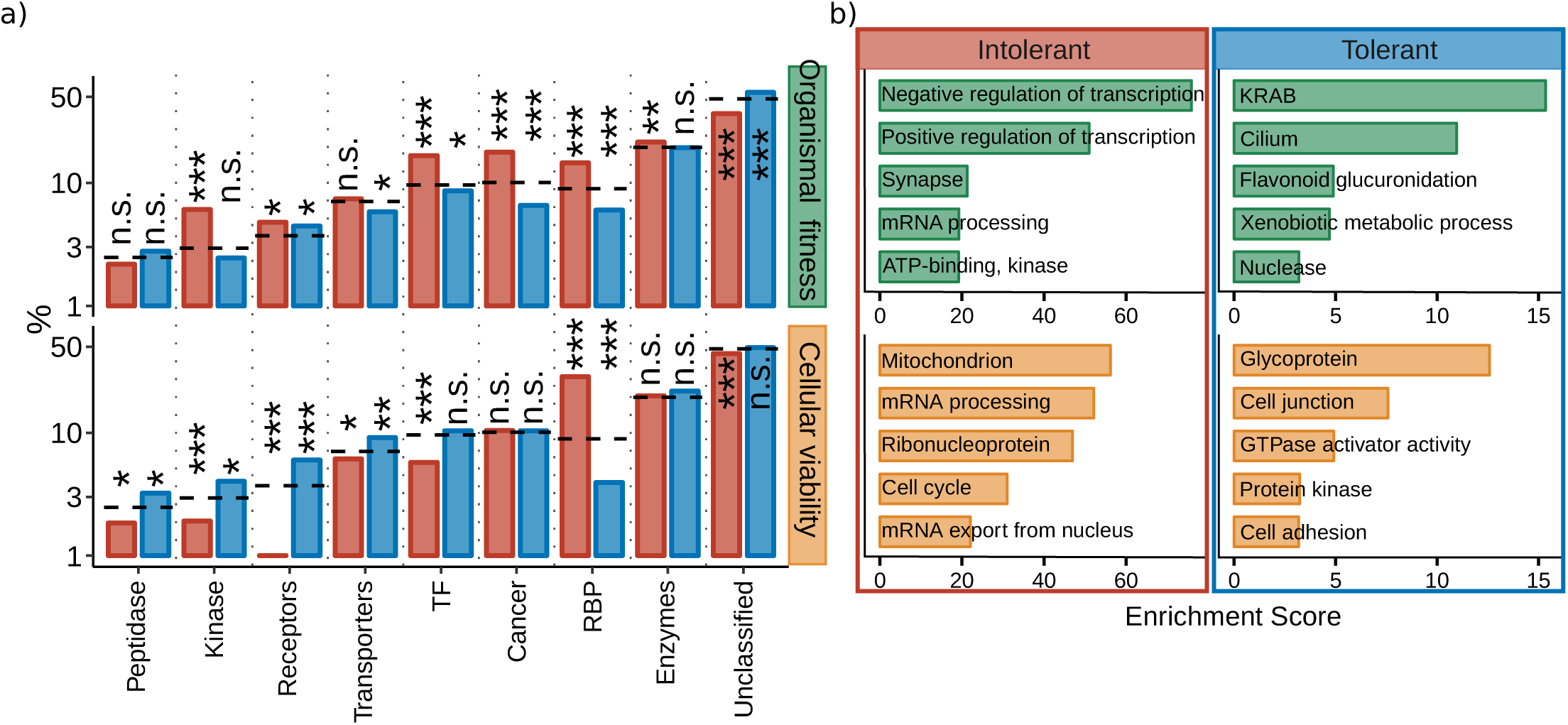
Protein class distribution and gene ontology term enrichment. **a**, Percent of genes belonging to each protein class distribution, dashed horizontal lines indicate the random expectation, deviation from expectation is shown on top of each bar. (significance: *p* < 0.05 = n.s., 0.001<*p*<0.05 = *, 0.0001<*p*<0.001 = **, *p*<0.0001 = ***) **b**, Top five enriched terms from functional cluster enrichment (DAVID) for each gene set.

### Molecular classes predict context-dependent mutational tolerance

Our previous results revealed differences in the evolutionary history of tolerance gene classes that affect CV or OF, suggesting that the deep conservation of intolerant genes with effect in cellular viability possibly stems from their role in core unicellular functions. To further explore this inference and unravel the differences between the OF and CV gene sets, we analysed the differential overrepresentation of tolerance gene groups within protein classes and gene ontology terms.

We again found differences in the protein class distribution of (in)tolerant genes, depending on the context of influence CV or OF. (Figure 5a). Protein kinases, receptors, transcription factors (TF), and cancer associated proteins show an unexpected contrasting pattern in CV vs OF context, with enrichment of OF intolerant genes but depletion of CV intolerant genes, and a reverse pattern for tolerant genes: enrichment for CV and depletion for OF. Thus these four protein categories, which together have a key role in developmental processes and associated signaling pathways, tend as a group to not tolerate LoF mutations in the human population, yet are not strongly required for human cell viability.

Gene ontology enrichment analysis further supports the difference between tolerance groups depending on CV or OF context, with distinct over-represented term (Figure 5b). Consistent with the previous result, OF intolerant genes show over-representation of gene ontology terms related with development and cell communication, such as transcription regulation, kinases, and synapse. CV intolerant genes on the contrary, show over-representation of core cellular processes related to cell energetics, replication, transcription, and translation. Consistent with contrasting functional properties of mutational gene tolerance classes depending on context, human genes that tolerate LoF mutations in cell culture are over-represented in processes related to cell adhesion and communication -- i.e., in similar processes that, in sharp contrast, do not tolerate mutations in the human population context (OF) (Figure 5b).

### Contrasting tissue-specificity and developmental activity of (in)tolerant genes

While global network structural and molecular functional properties provide evidence of consistent strong functional constraint on genes that do not tolerate LoF mutations in either human populations (OF context) or in human culture experiments (CV context), more detailed analyses of evolutionary history, protein classes, and gene ontology terms suggest that the two contexts (OF and CV) capture distinct functional roles of intolerant genes in the organism. To further explore the hypothesis of contrasting functional constraints, we gathered and interrogated data informative of developmental involvement and tissue-specific expression.

First, we evaluated the distribution of developmental stages in which genes are first expressed. Intolerant genes are expressed earlier in development than tolerant genes in both OF and CV contexts (Figure 6a), with at least 98% of the genes already expressed in prenatal stages. On the contrary, tolerant genes are depleted in prenatal stages and preferentially expressed after birth. This similar pattern of early expression is consistent with the high involvement of both TF mediated specification, cell-attachment, and core cellular replication in embryogenesis and organogenesis. However, when considering curated gene sets involved in specific developmental processes, we found a contrasting pattern between OF and CV gene sets, consistent with previous results. In OF context, intolerant genes are over-represented in every developmental category, while tolerant genes are depleted in all categories. In sharp contrast, in CV context, tolerant genes are over-represented in developmental processes, while intolerant genes are depleted (Figure 6b). This result further supports the view that genes intolerant of LoF mutations in the human population are preferentially involved in organismal development, and that the same constraint is not captured by *in vitro* screens of gene essentiality.

**Figure 6.**
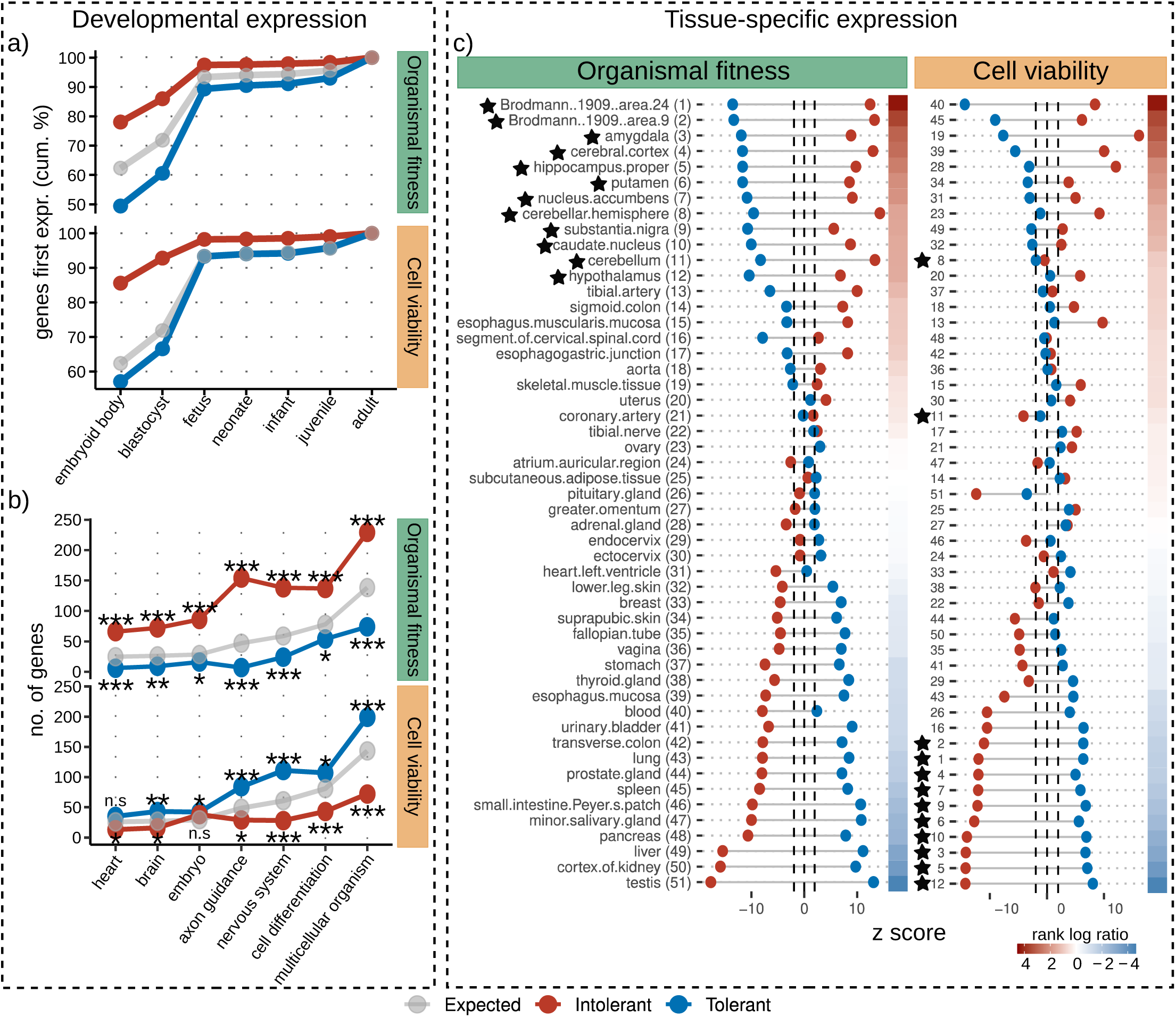
Gene set enrichment in tissue-specific expression, temporal stage expression, and developmental processes. **a**, Cumulative distribution of gene set percentage of genes first expressed by developmental stage. **b**, Number of genes per gene set associated with a developmental process. **c**, Expression enrichment deviation by tissue, tissues are ordered according to rank log ratio. Tissues highlighted with a star are part of the central nervous system. (significance: *p* < 0.05 = n.s., 0.001<*p*<0.05 = *, 0.0001<*p*<0.001 = **, *p*<0.0001 = ***).

In addition to developmental-stage associations, we next explored whether (in)tolerant gene classes of CV or OF context recover distinct preferential behavior in adult tissues. We used RNA-seq data from the Genotype-Tissue Expression (GTEx) project ^22^ to analyze patterns of tissue-specific expression. First we performed gene expression specificity analysis to compute for each tissue and gene a quantitative measure of specific expression relative to all other tissues (Methods). Then, we used the specificity values to estimate the degree to which a gene tolerance group shows unexpectedly high preferential expression in a given tissue relative to random expectation, resulting a z-score. We performed these calculations independently for each tissue and tolerance group, and for each context (CV or OF). We again found both common and particular patterns of behavior among OF and CV contexts. We found consistent opposite behavior in tissue preference in tolerant vs intolerant genes: in both contexts tissues with preferential expression of intolerant genes show depleted preferential expression of tolerant genes, and vice versa (Figure 6c).

Next, to contrast the tissue-preference behavior of tolerance groups in each context, we ordered the tissues by their relative preference to preferentially express intolerant vs tolerant genes. We measured this by the log ratio of the rank of the tissue in intolerant preferential expression relative to its rank in tolerant preferential expression. Using this approach, tissues that tend to preferentially express intolerant genes and not tolerant genes appear on top (Figure 6c). Notably, this analysis uncovered a contrasting behavior between OF and CV contexts: tissues from the central nervous system as a group (n=12 brain regions) show the largest relative preference of intolerant gene expression in the OF context and the least preference in the CV context. In other words, we found that the adult human brain tends to preferentially express genes that do not tolerate LoF mutations in the human population and to not preferentially express both OF tolerant genes and genes required for cell viability (CV intolerant genes) (Figure 6c). The other tissues do not show any clear pattern distinguishing CV and OF measures. We corroborated the reproducibility of these results by using independent gene expression reference data form human protein atlas^23^ (Supplementary Figure 3).

### Genes with inconsistent mutational tolerance behavior

The contrasting patterns found in OF versus CV gene groups suggests that genes with an inconsistent tolerance behavior across contexts might be driving the observed functional differences. To test this hypothesis, and to identify specific genes that capture the differential functional constraints accessible through population-based vs CRISPR-bases essentiality estimations, we defined consistency gene classes and repeated all association analyses, using this time only the new groups (Fig. 7a). We identified a group of 567 genes that do not tolerate LoF mutations in human populations, but that are not required for survival in human cells (organismal intolerant but cellular tolerant genes, OI-CT). Similarly, we identified a group of 351 genes that are cellular intolerant but organismal tolerant (CI-OT). Consistent with our previous results, OI-CT genes include major TF regulators of early cell lineage specification (e.g., SOX1, PAX6, and OLIG1), members of signaling pathways regulating these TFs (NOTCH1, NOTCH3, SMAD1), and genes encoding proteins relevant for non-cell autonomous physiological integration (e.g. ion channels CACNA1C, CLCN3, GABRA1) (Supplementary Table 1).

**Figure 7.**
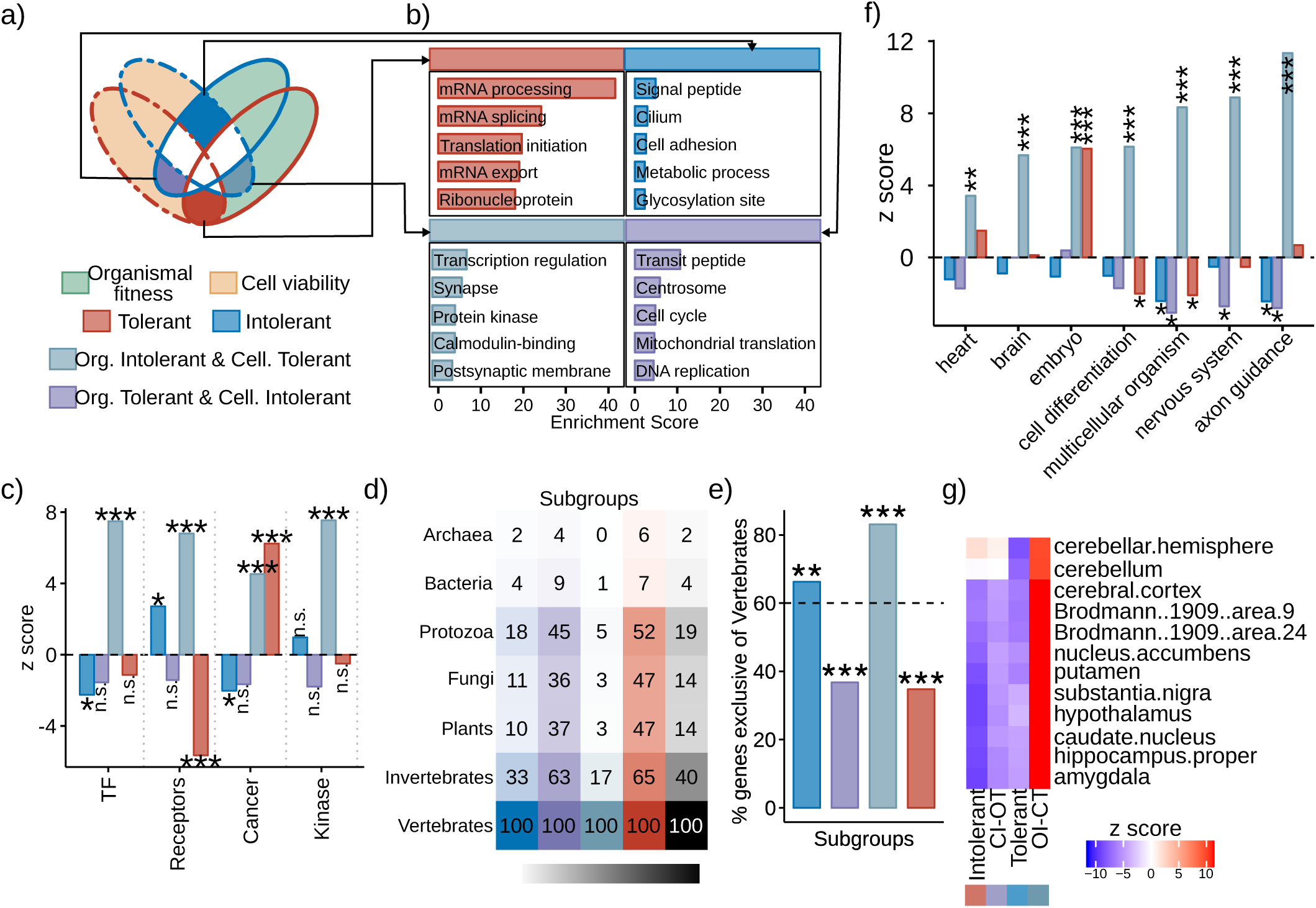
Consistent and inconsistent subgroups analyses. **a**, Subgroups definitions based on the overlap between OF and CV gene sets. **b**, Ontology term enrichment for each subgroup. **c**, Protein class distribution enrichment. **d**, Percent of orthologs in each taxon, color of the column indicates subgroup, black column shows the expected values. **e**, Percent of genes that only have orthologs among vertebrates. **f**, Enrichment of gene presence in developmental processes. **g**, Brain tissues expression enrichment. (significance: *p* < 0.05 = n.s., 0.001<*p*<0.05 = *, 0.0001<*p*<0.001 = **, *p*<0.0001 = ***).

As expected, association analyses revealed clear differences in these two inconsistent groups (OI-CT and CI-OT), in particular with respect to categories with contrasting behavior in OF versus CV gene sets. OI-CT genes are associated with gene ontology terms related to transcriptional regulation and neuronal communication, while CI-OT genes are associated with unicellular functions (Fig. 7b). Similarly, protein classes with contrasting behavior in OF vs CV (i.e., TFs, receptors, and kinases) are highly over-represented in OI-CT (Figure 7c). Notably, the same enrichment patterns are not observed when considering (in)tolerant genes with consistent behavior in both OF and CV contexts.

We similarly identified discrepancies in the evolutionary history of the new gene groups. OI-CT genes have less orthologs than expected within every taxonomic group except for Vertebrates, suggesting that many of these genes emerged relatively late in evolution, in pair with the emergence of Vertebrates (Figure 7d). To further explore this observation, we calculated the number of genes in each tolerance gene subgroup that have one-to-one orthologs only within Vertebrates, and not in other taxonomic groups. This analysis confirmed that more than 80% of OI-CT genes are exclusive to the vertebrate branch, in sharp contrast with both consistently intolerant genes and with CI-OT genes (Figure 7e). Lastly, OI-CT genes are also highly over-represented within every developmental process considered (Fig. 7f) and are preferentially expressed in adult human brain tissues (Fig. 7g). The evolutionary and functional patterns associated with the OI-CT group suggests that the genes in this group are relevant for organismal physiology.

Altogether, these results indicate that the genes with inconsistent mutational tolerance behavior across OF and CV contexts drive the differences observed in OF versus CV gene groups. That is, because specific genes do not tolerate LoF mutations in the human population but are not required for human cell viability, and, in contrast, other genes are required for the latter but do tolerate LoF mutations across the population; in vivo or in vitro overall estimations of human gene essentiality capture contrasting functional constraints overall.

## Discussion

We examined the degree to which measures that rank human genes according to their degree of tolerance to LoF mutations capture functional constraints. We considered tolerance estimations based on either in vivo exome-based population data or in vitro CRISPR-based perturbation experiments. To interpret evidence of functional constraint, we integrated genome-wide datasets related to gene function; including structural, functional, and evolutionary features; and compared essential genes versus genes with extremely high mutational tolerance, considering random proteome expectation. Our results indicate that intolerant genes (1) form a core network neighborhood in the human protein interactome, (2) are enriched in molecular properties suggestive of functional constraint, (3) are evolutionarily conserved, and (4) show preferential expression in specific tissues and developmental stages (Figure 8). The molecular and network properties that consistently discriminate intolerant from tolerant genes suggest that essentiality estimates based on mutational tolerance inference do recover functional constraints, irrespective of estimation context (OF or CV). However, we also found differences in the discriminatory properties of genes depending on whether their tolerance to mutation was estimated at the organismal or cellular level.

**Figure 8.**
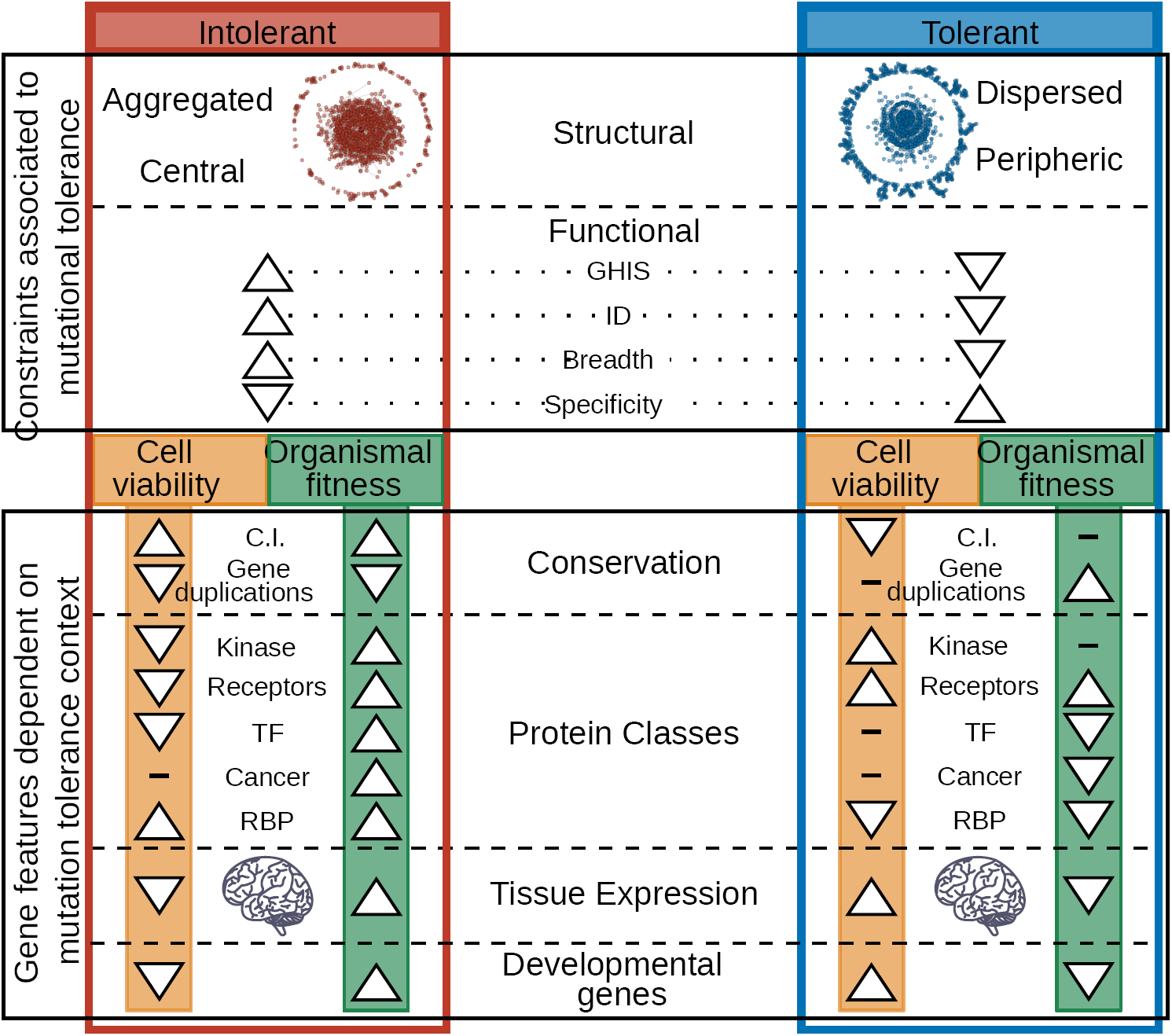
Results summary. Enrichment patterns of the main features found associated to genes essentiality. Top panel: Structural and functional features distinctive of tolerant/intolerant genes irrespective of the context. Bottom panel: Distinctive features that show a divergent pattern depending on the context in which mutational tolerance is defined.

Consistent with previous observations ^6,14^, we found that structural network properties consistently discriminate tolerance/essentiality classes. Intolerant genes are central and localized in the human interactome, while tolerant genes are dispersed in the periphery. This relative organization predicts a preferential vulnerability of the cell to intolerant gene failure, a principle that we demonstrated here by simulated network perturbation analysis. Both centrality and perturbation results provided results consistent with the hypothesis of a dominant role of intolerant genes in influencing cell behavior. Molecularly, intolerant genes also show properties often associated with gene functional relevance. We observe a significant tendency (FDR<0.01) for intolerant genes to be haploinsufficient, to have structural disorder, and to be broadly expressed. In contrast, tolerant genes show underrepresentation of these properties. Independent observations that further support relevance for cell behavior: broadly expressed and intrinsically disordered and are highly pleiotropic, given their structural flexibility and interaction promiscuity ^24^, while perturbations that manifest in dosage alterations of haploinsufficient genes might similarly be detrimental.

From an evolutionary perspective, we also observe a clear distinction between tolerant and intolerant genes, consistent with previous reports ^19,21^. In particular comparing tolerance classes, genes intolerant to LoF mutations have an older evolutionary history, with more one-to-one orthologs across species. Notably, we found that this evolutionary pattern is accentuated in intolerant genes estimated at the cellular-level compared with those measured at the organismal-level. This suggests that CV intolerant genes are more prone to be involved in basic cell-autonomous processes shared among all taxa, with prominent presence in unicellular organisms. In contrast, OF intolerant genes are either not enriched in unicellular groups (Archaea and Protozoa) or are underrepresented in Bacteria, suggesting that these genes emerged more recently in evolution and acquired a central role at a higher level of organization.

The idea that the context at which mutational tolerance is estimatimated discriminates genes operating at different levels of organization is reinforced when considering the differences we found for the enrichment of molecular classes among tolerance groups. The ontology terms associated with CV intolerant genes (mitochondria, RNA processing, ribonucleoprotein and cell cycle) relate to core functions required for cell survival, such as cellular metabolism and replication. On the contrary, CV tolerant genes relate to intercellular adhesion and communication (glycoprotein, cell junction, protein kinase). In extreme contrast with the cellular level, OF intolerant genes are enriched in functional features key to multicellularity, such as organismal development and cell-cell communication; as evidenced by over-representation of transcriptional regulators, synapse genes, and proteins of class kinase, receptor, and transcription factor. These results suggest that OF measures recover functional constraints stemming from multicellularity and organismal regulation, a property not readily captured by CV estimations. Consistent with this view, by considering curated gene annotations for developmental processes, we found a contrasting pattern between CV and OF genes. Genes involved in developmental processes are enriched for OF intolerant genes and CV tolerant genes, and depleted in OF tolerant genes and CV intolerant genes.

This contrasting behavior is also present in the tissue-specificity of gene expression. Specifically, we found that OF intolerant and CV tolerant genes are preferentially expressed in the adult human brain, in contrast with the underexpression of both OF tolerant genes and genes required for cell viability (CV intolerant genes). The overrepresentation of OF intolerant genes in the adult brain requires a careful explanation on their specific function in the organism and why is their sequence so mutationally constrained, which is beyond the scope of this paper. From the current results we speculate that, in addition to multicellular functional constraints, the depletion of LoF mutations estimated in human populations might capture constraints stemming from functional properties of species-specific relevance, such as higher cognition and associated traits grounded on the complexified human brain ^25^.

Although essentiality estimates from both cellular and organismal contexts do recover functional constraints, we found that a subgroup of 567 genes estimated as essential at the organismal level, yet nonessential at the cellular level, is responsible for the contrasting functional patterns found between OF and CV intolerant genes. These genes, which we refer to as (*OrgEssential*), are enriched in developmental processes, transcriptional regulation, and neuronal communication; and are preferentially expressed in the human brain. Despite being evolutionary younger than other essential genes, sharing one-to-one orthologs mainly with Vertebrates, *OrgEssential* genes seem to have developed a central role in the organism, providing an example of how during evolution novel genes can acquire essentiality properties by acting at levels of biological organization beyond core cell functionality.

## Methods

### Gene essentiality

Human gene essentiality estimations based on measures of tolerance to LoF mutations were taken from ^6^. These include the following scores based on the Exome Aggregation Consortium (ExAC) sample of 60,706 human exomes ^14^: residual variation intolerance score (RVIS) ^26^, EvoTol ^27^, missense Z-score ^28^, LoFtool ^29^, probability of haploinsufficiency (Phi) ^30^, probability of loss-of-function intolerance (pLI) ^14^, and selection coefficient against heterozygous loss-of-function (shet) ^31^; and scores based on cell culture perturbation-based experiments performed in KBM7, Raji, Jiyoye, HCT116, and K562 cell lines in ^12^; the KBM7 cell line in ^10^, and RPE1, GBM514, HeLa, and DLD1 cell lines in ^32^.

### Intrinsic Structural Disorder

Disorder predictions for each protein in the human proteome were generated at residue resolution using IUPred ^33^. A gene intrinsic disorder score was calculated by averaging the predicted residue scores over the corresponding protein. Scores range from 0 to 1, with higher scores indicating a higher propensity toward intrinsic disorder.

### Haploinsufficiency

A predictive genome-wide haploinsufficiency score (GHIS) was obtained from ^34^.

### Gene expression specificity

Reference RNA-seq data for human tissues was downloaded from the Genotype-Tissue Expression project (GTEx.v7) ^22^ (https://www.ebi.ac.uk/arrayexpress/experiments/E-MTAB-5214/), and the Human protein atlas (HPA) ^23^ (https://www.ebi.ac.uk/arrayexpress/experiments/E-MTAB-2836/). The GTEx dataset includes 53 tissues profiled from 961 donors. The HPA dataset includes 32 tissues profiled from 122 control subjects. For both datasets the median expression over replicates was considered as the expression value of the tissue. Expression breath values for each gene were calculated as the fraction of tissues in which the gene is expressed, using an arbitrary cut-off value of 2 RPKM to determine expression. Expression specificity was measured using the Tau statistic on the same tissue-median matrix as computed in ^35^.

### Protein classifications

Proteins were classified as Transcription factors (TF), Transporters, Receptors, Enzymes, Peptidase, Kinase, Cancer-related, and RNA binding proteins (RBP) based on combined curated annotations extracted from the Human Protein Atlas (https://www.proteinatlas.org/humanproteome/proteinclasses) ^23^, TF reference in ^36^, RBPs reference from ^37^, and transporters and receptors reported in ^38^.

### Evolutionary conservation

Comprehensive gene homology information for each human gene with respect to 187 species was extracted from Ensembl comparative genomics resources ^39^. Only one-to-one orthology relationships were considered to build a binary gene-species matrix. A gene conservation index was calculated for each human gene as the fraction of species having a corresponding ortholog ^39^. Gene duplications data was extracted from ^40,41^.

### Developmental annotations

Developmental expression classes and developmental process gene annotations were downloaded from the Online Gene Essentiality database (OGEE.v2) at (http://ogee.medgenius.info/downloads/) ^41^.

### Mutational tolerance gene group definition

We define consensus organismal fitness and cell viability mutational tolerant and intolerant groups by first independently calculating the gene average rank across n=6 measures of *in vivo* tolerance to LoF mutation for OF context, and n=10 *in vitro* measures for CV context. In all measures, as reported in ^6^, values increase with the degree of intolerance to mutation: intolerant genes have high values. The group of Tolerant genes was defined as the bottom 20% genes with consensus score rank (lowest constraint), and Intolerant genes as the top 20% (highest constraint). 4 additional subgroups were defined based on the overlaps between intolerant and intolerant groups (Figure 2c). The resulting subgroups are: consistent tolerant genes (n=714 genes classified as tolerant in both conditions), consistent intolerant genes (n=771 genes classified as intolerant in both conditions), organismal intolerant but cellular tolerant genes (OI-CT) (n=567 genes classified as OF intolerant and CV tolerant), and cellular intolerant but organismal tolerant genes (CI-OT) (n=351 genes classified as OF tolerant and CV intolerant).

### Analysis of gene set aggregation and centrality in the PPI network

A reference human protein-protein interaction (PPI) network was obtained from ^42^.

Gene set aggregation in the PPI network was quantified using three complementary approaches: module size of the gene set subgraph, clustering enrichment and pairwise distance distribution. Subgraph module size was calculated by counting the number of nodes (Sc) and edges (Cc) of the largest connected subgraph formed by proteins belonging to a given gene set. Clustering enrichment was measured using the SANTA method ^43^. Pairwise shortest distance between every protein pair was measured using the *igraph* R package ^44^. The distance distribution of each gene set, was characterized by calculating its minimum (*D*_*s*_) and mean (*D*_*sm*_) distances. Network centrality was calculated using three complementary measures: degree, betweenness, and coreness. These measures were quantified for every node using the *igraph* R package ^44^. For each gene set, network aggregation and centrality enrichment was calculated comparing the gene set mean with a random distribution obtained from 10,000 randomly sampled gene sets of the same size, resulting in a z score.

### Network structural robustness analysis

Network robustness was characterized by measuring the effect on network structure after the targeted removal of nodes according to the OF and CV mutational tolerance ranking, and comparing it with random expectation. Network structural response was assessed calculating the number of nodes (*S*_*f*_) and edges (*C*_*f*_) in the perturbed giant connected component after removing a fraction (*f*) of nodes relative to the unperturbed measures. For each measure, we built a random expectation by removing fractions from 0.01 to 0.99 of randomly selected nodes, repeating this procedure 10,000 times.

### Enrichment of gene set functional properties and protein class distribution

Gene set enrichment in gene functional features (GHIS, ID, expression specificity, expression breadth, earliest stage expression, and developmental process annotation) was calculated by comparing the gene set mean with the random expectation obtained from measuring the given feature in 10,000 randomly sampled gene sets of the same size, resulting in a z score. Deviation in protein class distribution among gene sets was calculated by determining the percentage of genes belonging to each protein class and comparing it to the random expectation built from 10,000 randomly sampled gene sets of the same size, resulting in a z score.

### Gene set evolutionary analysis

For each gene set, deviation of conservation index and number of gene duplications were calculated by a z test with a random distribution obtained from 10,000 randomly sampled gene sets of the same size. The gene-species ortholog matrix was reduced by classifying species by taxonomic group resulting in: Archaea (21 species), Bacteria (99 species), Protozoa (16 species), Fungi (8 species), Plants (9 species), Invertebrates (24 species), and Vertebrates (10 species). Using the reduced orthologs matrix, the percentage of orthologs in each taxonomic group was calculated for each gene set.

### Tissue specific and expression enrichment

We used RNA-seq data from the Genotype-Tissue Expression Project (GTEx) (51 tissues) ^22^ and from the Human Protein Atlas (HPA) (32 tissues) ^23^ to measure tissue specific differential expression of every gene. Gene differential expression was calculated using *voom* and *lmFit* functions from the *limma* package in R ^45^. With the genes-tissue differential expression matrix, we measured the degree of preferential enrichment of each gene tolerance group in every tissue. To do this we compared the mean gene differential expression of every group to random expectation, calculated from 10,000 randomly sampled gene sets, and computed the differential expression deviation (z score) of each tolerance group in every tissue.

### Gene set enrichment analysis

Gene ontology enrichment analysis was performed using DAVID (https://david.ncifcrf.gov/summary.jsp) ^46,47^.

## Supplementary materials for

**Figure S1.**
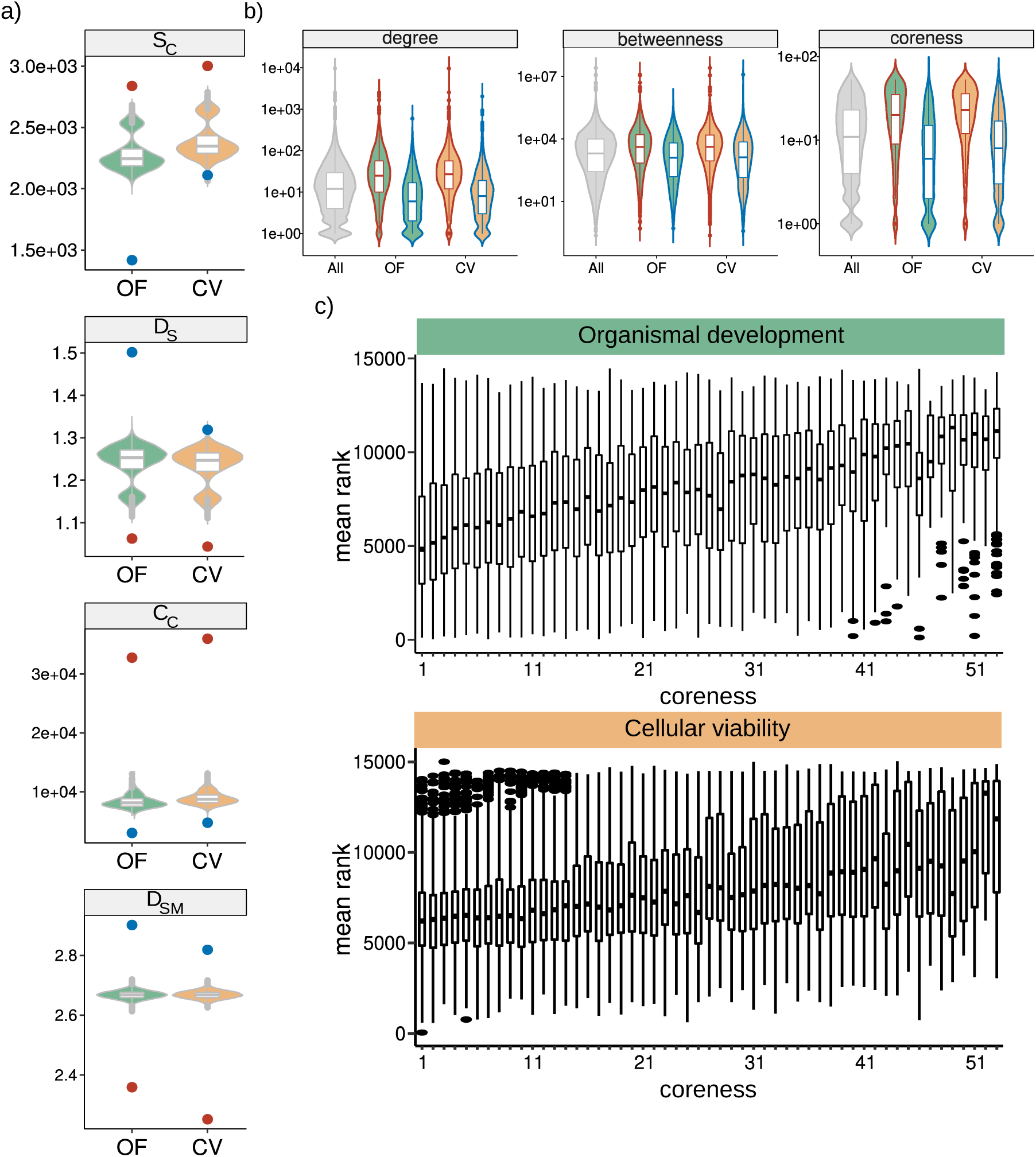
PPI structural features distribution. **a**, Aggregation measures: mean random distribution is shown as a violin diagram and the observed value for tolerant and intolerant gene sets as a dot colored corresponding to the group. **b**, Centrality measures: distribution of node centrality, line colors correspond to the tolerant/intolerant condition and filling color to the context. **c**, Boxplot showing the relationship between coreness and mutational tolerance.

**Figure S2.**
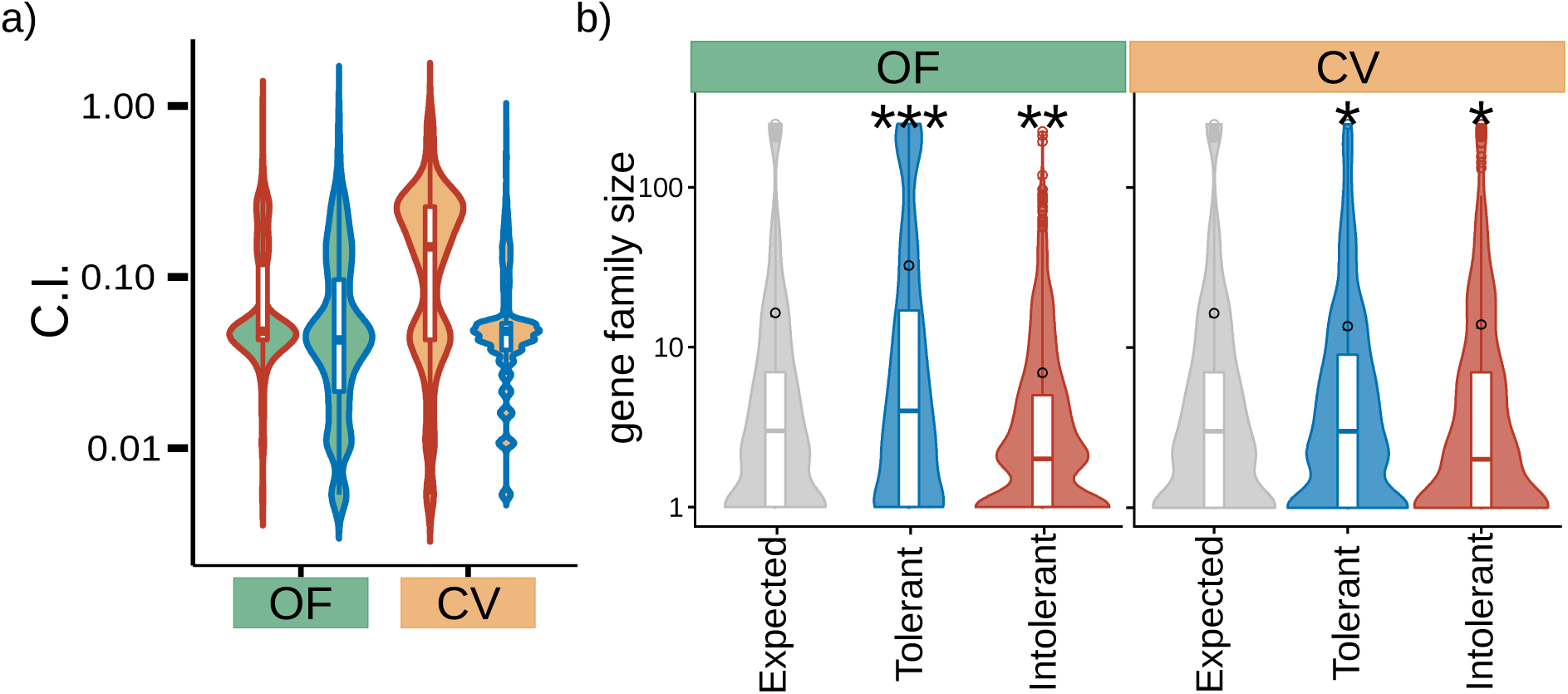
Evolutionary patterns. **a**, Gene conservation index distribution among gene sets. **b**, Distribution of the number of genes in a family for each gene set. Black circle represents the distribution mean.

**Figure S3.**
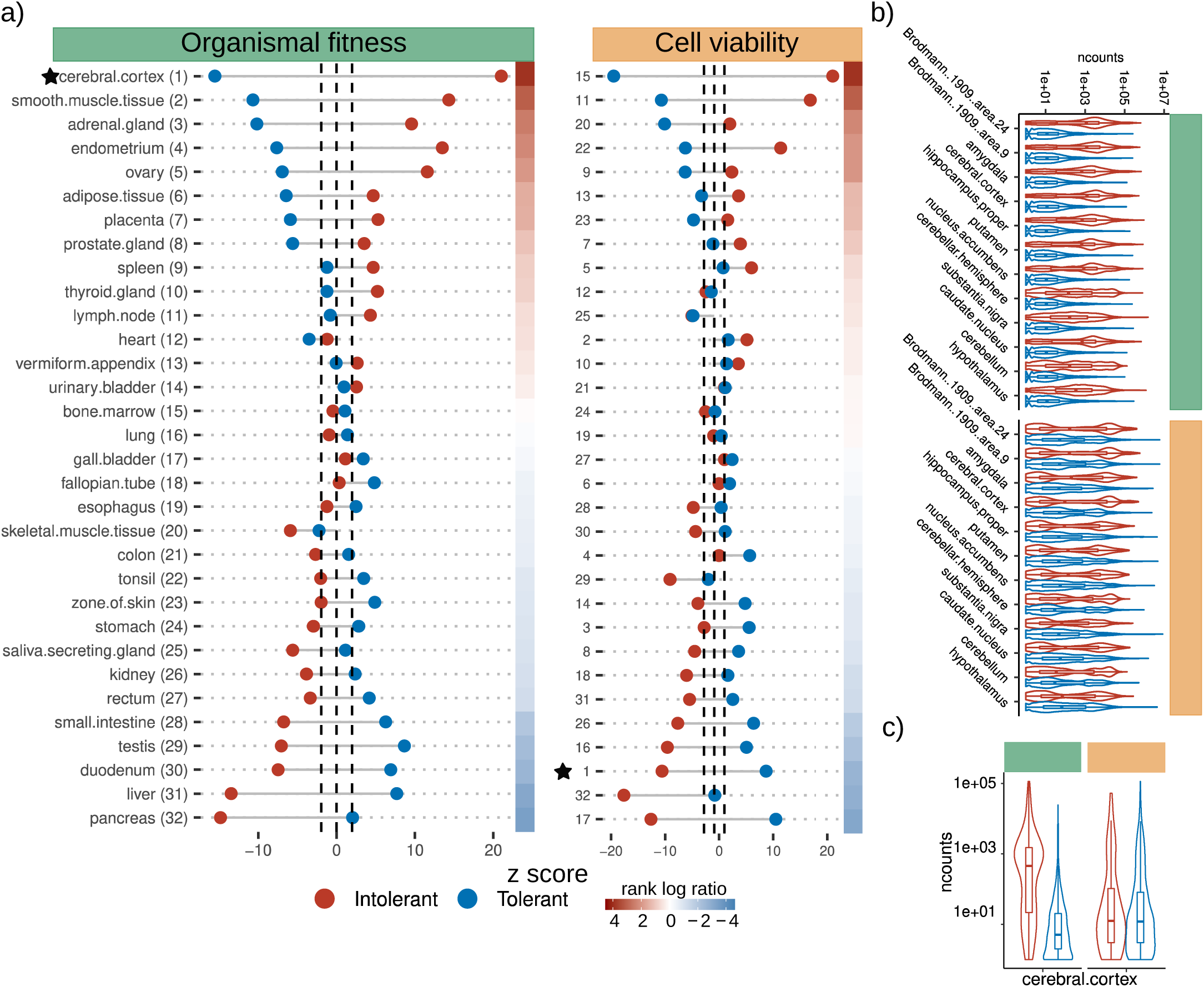
HPA tissue enrichment. **a**, Expression enrichment deviation by tissue calculated from HPA expression dataset. Each point represents the deviation in expression enrichment for the given gene set and condition in a particular tissue, tissues are ordered according to rank log ratio. Tissues highlighted with a star are part of the central nervous system. **b**, Distribution in the transcript number for each tissue from the central nervous system from GTEx data. **c**, Same as b) for HPA data.

## List of Tables

**File S1. Tables with genes mutational group membership and analyzed features scores**

**File S2. Genes one-to-one orthologs.**

